# Time-Optimal Adaptation in Metabolic Network Models

**DOI:** 10.1101/2022.02.05.479110

**Authors:** Markus A. Köbis, Alexander Bockmayr, Ralf Steuer

## Abstract

Analysis of metabolic models using constraint-based optimization has emerged as an important computational technique to elucidate and eventually predict cellular metabolism and growth. In this work, we introduce *time-optimal adaptation* (TOA), a new constraint-based modeling approach that allows us to evaluate the fastest possible adaptation to a pre-defined cellular state while fulfilling a given set of dynamic and static constraints. TOA falls into the mathematical problem class of *time-optimal control problems*, and, in its general form, can be applied in a broad sense and thereby extends most existing constraint-based modeling frameworks. Specifically, We introduce a general mathematical framework that captures many existing constraint-based methods and define TOA within this framework. We then exemplify TOA using a coarse-grained self-replicator model and demonstrate that TOA allows us to explain several well known experimental phenomena that are difficult to explore using existing constraint-based analysis methods. We show that TOA can explain accumulation of storage compounds in constant environments, as well as overshoot uptake metabolism after a period of nutrient scarcity. TOA reveals that organisms with internal temporal degrees of freedom, such as storage, can in most environments outperform organisms with a static intracellular composition. Furthermore, TOA shows that organisms adapted to better growth conditions than present in the environment (“optimists”) typically outperform organisms adapted to poorer growth conditions (“pessimists”).

## 2 Introduction

Over the last decades, several modeling frameworks have been proposed to understand the organization and functioning of cellular metabolism and growth. Some of the most popular approaches are constraint-based methods, in particular *flux-balance analysis* (FBA) [21]. Constraint-based methods typically make use of optimality principles that are motivated by evolutionary arguments. That is, instead of requiring a detailed mechanistic understanding of the underlying regulatory machinery, properties of cellular metabolism, such as exchange fluxes or biomass accumulation, are predicted based on the assumption that metabolism has evolved according to certain evolutionary optimality principles.

More recently, constraint-based methods have been extended to quantitatively account for the synthesis costs of the biological macromolecules that are required for cellular metabolism and growth, giving rise to *resource balance analysis* (RBA) [8] and integrated reconstructions of *Metabolism and macromolecular Expression* (ME) [15]. While the initial approaches were restricted to time-invariant environments and subject to steady-state conditions, various dynamic extensions have also been proposed, such as *dynamic FBA* (dFBA) [18], *dynamic enzyme-cost FBA* (deFBA) [30], *conditional FBA* (cFBA) [26, 25], *dynamic RBA* (dRBA) [13], *dynamic ME* [31], and *regulatory dynamic enzyme-cost FBA* (r-deFBA) [17]. These dynamic frameworks are computationally more expensive and allow predicting time courses over a given time interval, such that the variables fulfil a certain (linear) optimality principle. Typically, within these frameworks, the time intervals over which the solutions are considered are predefined.

In this work, we extend these existing approaches and propose *time-optimal adaptation* (TOA) as a new constraint-based modeling framework that allows us to evaluate the fastest possible adaptation to a pre-defined cellular state while fulfilling a given set of dynamic and static constraints. If the underlying dynamics of the biological system are governed by differential equations (ODEs) subject to algebraic constraints such as positivity, that is, so-called differential-algebraic equations (DAEs), time-optimal adaptation falls into the mathematical problem class of *time-optimal control problems*, which are optimal control problems where the time-interval is part of the objective [10]. In its general form, time-optimal adaptation can be applied in a very broad sense and thereby extends most of the existing constraint-based modeling frameworks.

Our approach allows us to compute feasible time courses to simulate or predict adaptations of cellular metabolism to environmental shifts. Potential applications include an analysis of cellular doubling, i.e., to analyze the optimal metabolic trajectory that results in a doubling of all cellular components in the shortest time, as well as an analysis of the temporal adaptation to changing nutrient availability.

We exemplify TOA using a coarse-grained self-replicator model [20, 6] and demonstrate that TOA allows us to explain several well-known experimental phenomena that are difficult to investigate using existing static or dynamic constraint-based analysis methods. In particular, we demonstrate that TOA can explain accumulation of storage compounds also in time-invariant environments–a counterintuitive fact that cannot be predicted using RBA and related methods. Likewise, we demonstrate that “luxury uptake” of nutrients, i.e., the fact that microorganisms may take up more of a limiting resource than strictly required for steady-state growth, can be explained by TOA and does not necessarily require competition within a microbial community. Furthermore, our analysis shows that organisms with internal temporal degrees of freedom, such as storage, can in most environments outperform organisms with a static intracellular composition. Finally, TOA shows that in constant (or slowly changing) environments, organisms adapted to better growth conditions (“optimists”) outperfom organisms adapted to poorer growth conditions (“pessimists”) when placed in the same environment.

## 3 Materials and Methods

### 3.1 Introduction and Notation

The dynamic simulation of metabolic networks by means of fully parameterized ODE/DAE models is an ideal scenario that, in most cases, cannot be met due to the inherent incompleteness and uncertainty of the description and the involved parameters. *Constraint-based modeling* [2] has therefore become an important paradigm for the computational description of cellular metabolism and growth. The general idea can be framed as follows: instead of making use of a fully mechanistic description of biochemical dependencies by means of reaction rate equations, the system is characterized by a set of constraints/inclusions, typically defined by (in-)equalities that constrain the dynamics over a time interval [*t*_0_, *t*_end_] of interest.

Before capturing our approach in mathematical terms in Section 3.2, we introduce some notation. See also Appendix A. The function 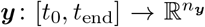 is used to describe the cellular dynamics by the *total amounts y*(*t*) of intracellular compounds at time *t* (typically measured in mol), with 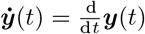 denoting the time-derivative. For simplicity, we focus on the dynamics of intracellular compounds only, extracellular compounds (e.g. nutrient or waste product concentrations) are not included in ***y***. Our framework, however, can be readily adapted to include the dynamics of extracellular compounds (see the Appendix B.3 for details). Furthermore, our description is based on the assumption of a well-stirred metabolism, i.e., the spatial distribution of compounds is not considered. We distinguish the total amounts of molecules *y*(*t*) from their *concentrations c*(*t*), defined by

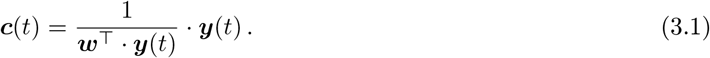

Here, the vector 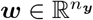 collects the molar masses of the entities of ***y*** such that

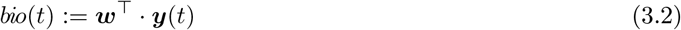

denotes the total biomass of the system.

The time evolution of the state vector ***y***(*t*) can be described by means of ordinary differential equations

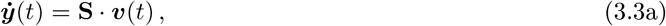

where 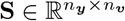 denotes the *stoichiometric matrix* and 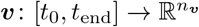 the *flux rates* of the reactions. Note that in (3.3a), the flux rates ***v***(*t*) may in general also depend on the environment the cell culture is exposed to.

Often, and specifically for large networks, the stoichiometric matrix 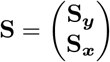 is split up such that ‘slow’ and ‘fast’ intracellular compounds are described separately, and (3.3a) is replaced by

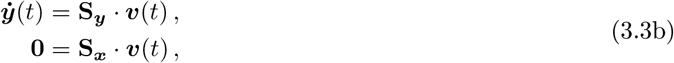

where the fast compounds, corresponding to the rows of **S_*x*_** are subject to a *quasi steady-state approximation* (QSSA). In this case, for simplicity of notation, the fast components will be removed from the vector ***y***(*t*).

### 3.2 Constraint-based Modeling

To capture the broad range of simulation frameworks that time-optimal adaptation is able to cover, we abstractly denote the constraints defining the specific constraint-based description of the cell via

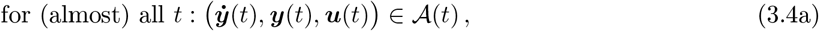

where the set 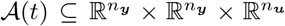 is typically defined through (in-)equalities such as steady-state assumptions and/or positivity requirements. The particular form of the set 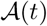 usually depends on the chosen modeling framework and its granularity. For the present work, we model the influence of the external conditions via the explicit time-dependence of 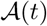. The vector-valued function 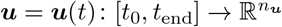 signifies the degrees-of-freedom of the cell, i.e., quantities that are not uniquely determined from the current state of the cell and its environment. In the context of control theory, ***u***(*t*) defines the controls; on the biochemical level, it can for example stand for flux rates ***v***(*t*) but also for parameters within the model.

[resume]The formal statement (3.4a) is usually not enough to sufficiently constrain the solutions, because the feasible region is too large to obtain biochemical insight. To get biochemically meaningful results, (3.4a) is therefore often accompanied by boundary conditions and an optimality principle, i.e., a global objective function *f* to be optimized:

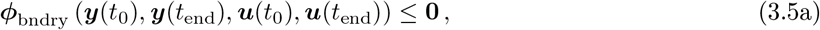

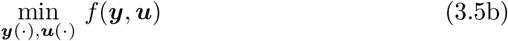

The boundary conditions (3.5a) are defined by means of inequalities to allow for more generality of this description. Usually, the boundary conditions will only contain initial values, provided by equality constraints, i.e., two inequalities. In some cases, optimality principles are already incorporated into the constraint set 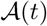, see the following examples.

In the context of optimal control-based methods with ODE/DAE constraints, the flux rates at any fixed point in time cannot (mathematically) be determined as they enter the problem as control variables [5]. This is why (3.4a) technically can only be enforced for *almost* all times. Numerically or with respect to the biochemical reasoning, however, this has no further implications. In the following, we illustrate how (3.4) provides an abstract framework to describe established examples of constraint-based modeling.

#### Example 3.1

(Dynamic FBA, dFBA). *Dynamic (or iterative) flux-balance analysis [29, 18], although one of the most commonly used dynamic frameworks within constraint-based modeling, is not consistently defined in the literature. Here, we refer to the formulation in [12], see also [11] for the characterization of dynamic FBA as a “dynamical system with a linear program embedded.”*

*The control quantities **u***(*t*) *can in this case be directly identified with the flux rates in the metabolic network model, i.e*., ***v***(*t*) = ***u***(*t*). *The overall dynamics are governed by* (3.3a), *positivity requirements on* ***y***(*t*) *and flux bounds **lb***, 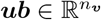:

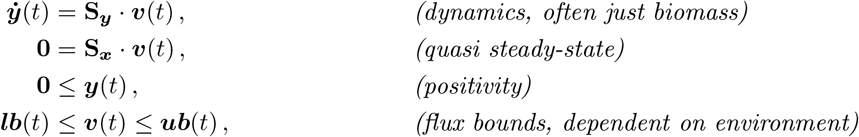

*with given initial conditions*

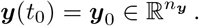

*The flux rates are determined through optimization of a linear functional (often the flux through the biomass reaction, assembled in a vector* 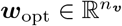)

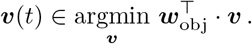

*The quantities in* (3.4) *can be identified as:*

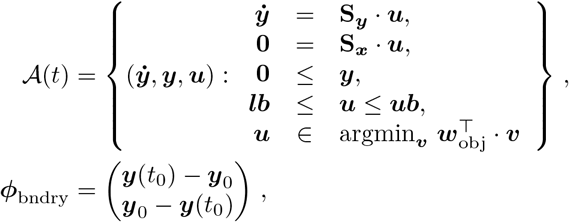

*while typically no additional (global) objective function is present. Note that the defining condition on the fluxes 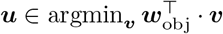 is an inclusion, such that the solutions to dynamic FBA problems are, in general, not unique. To remedy this*, flux variability analysis *(FVA, [19]) was introduced as a computational tool to explore the range of possible solutions of the static sub-problems*.

#### Example 3.2

(Dynamic enzyme-cost FBA, deFBA). *Dynamic enzyme-cost FBA [30] is a dynamic extension of FBA that takes into account the temporal development and function of the enzymes. This is modeled by a system of linear inequalities*

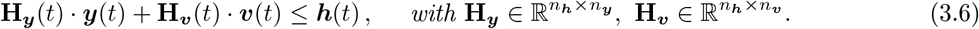

*The model is usually formulated as an initial-value problem*

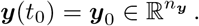

*Similar to FBA, deFBA assumes that a certain objective function is to be optimized. Since the framework entails a fully dynamic model over the whole time range of interest, the objective function contains ‘global’ information, expressed as an optimal control objective of Boltza-type [5]*,

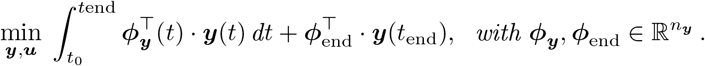

*Like in dFBA, the control variables in deFBA can be identified with the flux rates and the description in terms of* (3.4) *is given by*

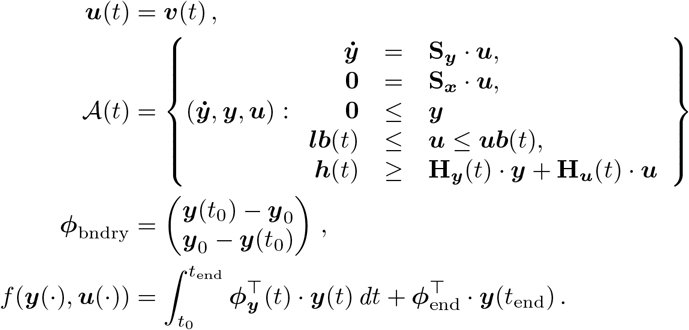

#### Example 3.3

(Conditional FBA, cFBA). *This framework [26, 25] is again a dynamic extension of resource balance analysis (RBA) [8]. Like in deFBA, enzymatic constraints (potentially alongside further constraints, e.g., on the cell’s density) are included via* (3.6). *The boundary values in cFBA, however, are defined through a periodicity condition that accounts for the growth of the cell:*

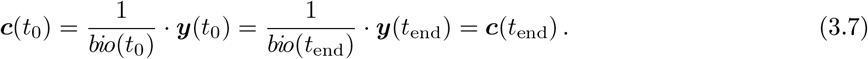

*Instead of using the biomass production on all time points, the objective in cFBA is the total growth of the cell until t*_end_. *In terms of* (3.4), *cFBA can be summarized as*

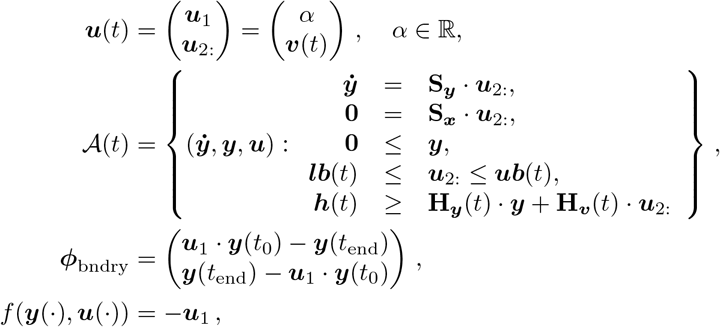

*where **u***_1_ *refers to the first component of the vector **u** and* ***u***_2:_ *to the vector of the remaining entries. If no constraints on the cell density are included in* (3.6), *the inequalities defining cFBA are often scale-invariant in the sense that for each solution **y***(*t*) *and each number β* ≥ 0, *the function β* · ***y***(*t*) *is also a solution. To exclude trivial solutions, the boundary conditions are therefore often extended such that the biomass at t*_0_ *is equal to one. Note that cFBA is inherently nonlinear as the products u*_1_ · ***y** in the boundary value constraints contribute quadratically in the unknowns **y** and **u**. Like in RBA, the numerical solution of cFBA problems therefore comprises a series of linear programs that have to be solved after a discretization of the dynamics by means of a collocation scheme*.

#### Example 3.4

(Iterative RBA, [16], see also dynamic ME [31]). *Just as dynamic FBA forms a dynamic extension of classical FBA by iteratively applying the algorithm with constraints following the external conditions, resource balance analysis (RBA, see [8]) can also be applied consecutively. In doing that, the limit case of infinitesimal short sub-intervals, leads to a fully dynamic framework. Numerically, this limiting process is skipped and one only solves RBA problems on a series of short—but finite—time intervals. Note that, as cFBA, RBA uses periodicity conditions like* (3.7) *which implies that, in constant external conditions, only one RBA problem needs to be solved. The full solution in this case is given by an exponential curve for **y***(*t*). *Note that there are fewer degrees-of-freedom for the cell when compared to deFBA, as the fixed concentration values for the metabolites in the case of iterative RBA also block internal dynamics of the metabolic network. A formal description in terms of constraint-based modeling as in* (3.4) *is provided in Appendix D*.

### 3.3 Time-optimal adaptation: definition and forms

Previous frameworks for constraint-based optimization did not explicitly include the time interval as part of the optimization objective. In the following, we introduce *Time-Optimal Adaptation* (TOA) as a framework to analyze transition between different cellular states in the shortest possible time. TOA is motivated by the assumption that under certain environmental conditions, cells may have evolved to reach a goal state ***y***^goal^ in the shortest possible time, starting from an initial state ***y***^init^. This transition might either take place in a variable environment, encoded by a time-dependent set 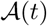, or in a constant environment. Likewise, the target amounts ***y***^goal^ may either have to fulfill additional optimality criteria, or may correspond to a pre-defined or experimentally measured state.

Mathematically, we capture such a strategy in the following way:

#### Time-Optimal Adaptation

Given an initial/current amount of molecules 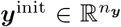 and a target amount 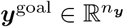, the optimization objective is to transition from the former to the latter as quickly as possible.

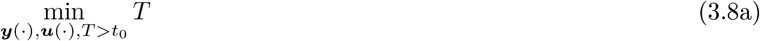

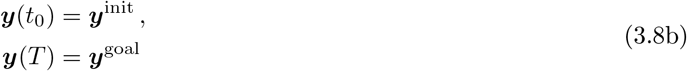

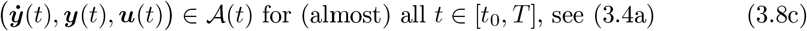

The constraints (3.8c) and (3.8b) can be framed within the abstract constraint-based framework (3.4) by including ***y***^init^ and ***y***^goal^ using

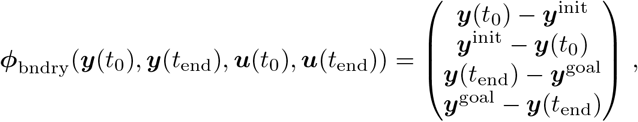

whereas the global objective function, cf. *f* in (3.4), does not explicitly contain any of the variables ***y*** or ***u***. Instead, the general framework of constraint-based modeling (3.4) is extended through time-optimal adaptation by using *the end point of the time interval of interest itself* as the optimization objective function. In contrast to the frameworks with non-time-dependent objective function as defined in (3.5b), TOA provides solutions (***y***(*t*), ***u***(*t*)) only on the time interval [*t*_0_, *T*] instead of (arbitrary) [*t*_0_, *t*_end_].

##### Remark 3.5.

Within this work, we use the term “adaptation” in a control-theoretic sense. That is, the term refers to changes in the intracellular amounts or concentrations in response to the environmental conditions, respecting the given constraints. In an evolutionary context, such changes are typically considered as “acclimation”.

##### Remark 3.6.

We do not require the constraint set 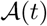 in the above formulation to have any specific form. This means that time-optimal adaptation can be defined irrespective of the concrete modeling paradigm underneath the simulation. Practically, even discrete time/state systems fit well within TOA. To be concise, however, in the following we concentrate below on frameworks closely related to deFBA and cFBA.

### 3.4 Applications and Case Studies

Next we introduce two particularly relevant applications of TOA.

#### Application 3.1

(Cell Doubling). *A first natural application of TOA is cell doubling, where the objective is to double all cellular components in minimal time, such that*

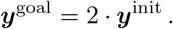

*The resulting trajectory thus can be interpreted as one cell cycle. Neither the initial, nor the goal amount have to be optimal with regard to other objectives. Within the TOA framework, cell doubling can be considered either in a constant environment, or with time-dependent external conditions*.

#### Remark 3.7.

Constraint-based optimization of metabolism typically does not distinguish between solutions for a single cell versus solutions for a homogeneous population of cells. Likewise, the time courses for cell doubling predicted by TOA can either be interpreted for a single cell or a homogeneous, synchronized population of cells. To describe cell doubling within a population of non-synchronous cells, that is, each cell within the population is at a different time point with respect to the progress through its cell cycle, we have to average over the population or, equivalently, over a full cell cycle. In Section 4.4, we consider the average cellular composition that is obtained from *in silico* measurements of a population of non-synchronous cells by averaging the respective concentrations over one cell cycle.

#### Application 3.2

(Transitions at a nutrient shift and the role of biomass). *A second important application of TOA is to consider a sudden change in the external conditions, i.e., from a given constant nutrient availability for t* < 0 *to a different one for t* ≥ 0. *In this scenario, TOA can be utilized to predict the transition of the intracellular amounts **y***^init^ *to new target amounts **y***^goal^. *The new target amounts might either be optimal with respect to the new environmental conditions (as defined by RBA), or be provided otherwise (for example by experimental observations). In both scenarios, the target amounts are typically defined in terms of concentrations instead of absolute amounts. Hence, we must also formulate the boundary conditions in terms of **c***(*t*),

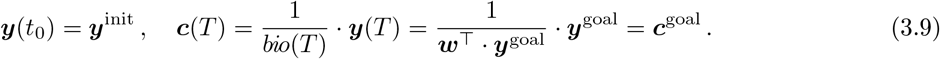

*As shown in Appendix C, it is possible to rearrange the conditions* (3.9) *such that a linear equality system in the unknowns* (***y***(*t*_0_), ***y***(*T*)) *is obtained. Therefore, the concentration-based definition has no immediate drawbacks regarding the numerical solution*.

*However, an as-quick-as-possible transition from one intracellular concentration to another does not incorporate the overall (i.e. biomass-) growth of the cell and thus might represent an evolutionarily implausible strategy. Rather, the transition to new external conditions involves a balance between fast transition to a (better adapted) novel state and the requirement to still increase the total biomass of the cell. We therefore propose a* two-objective optimization problem:

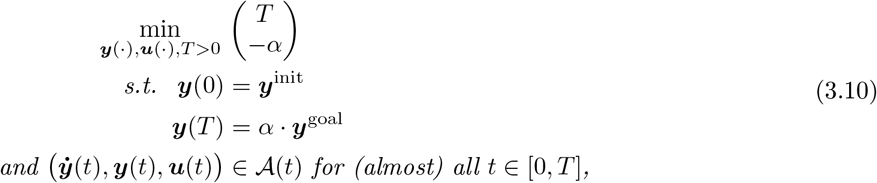

*where **y***^init^ *denotes a normalized vector of intracellular amounts which describe the cells for the environment t* < 0. *The “normalization” here can, for example, be understood as **w***^┬^ · ***y***^init^ = 1. *Accordingly, **y***^goal^ *denotes a normalized vector for the environmental conditions after the nutrient shift. ‘Minimality’ in* (3.10) *is to be understood in the sense of Pareto: a triple* (***y***(*t*), ***u***(*t*), *T*) *is optimal if T cannot be decreased without decreasing α such that **y***(*T*) = *α* · ***y***^goal^, *and vice-versa if α cannot be increased without also increasing the end time T. The set of all optimal solutions of* (3.10) *describes the different compromises between fast adaptation and continued growth*.

#### Remark 3.8.

Note that the boundary conditions (3.9) do not entail any direct condition concerning *bio*(*t*_end_) = ***w***^┬^ · ***y***(*t*_end_). If the metabolic network allows for a quick degradation of compounds, it might be optimal (in the sense of TOA) to drastically shrink before actually adapting to the new concentrations, or even to completely disintegrate all metabolic compounds to zero. Such a behavior would be in line with the description of time-optimal time courses as induced by (3.9). To remedy this, a linear inequality

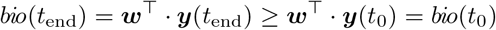

can be added, illustrating again the power of constraint-based modeling. Whenever necessary, this was done for the *in silico* experiments in Section 4.

### 3.5 Numerical Solution

The optimization problem (3.8) of TOA contains a very general condition on the dynamics of ***y*** in the form of Eqn. (3.8c). To design an algorithm able to cope with this generality, we assume that a numerical method is available that can simulate this dynamic behavior subject to boundary conditions on a given fixed time-interval [*t*_1_, *t*_2_] ⊆ [*t*_0_, *T*] and/or to determine whether such a solution exists. Provided this condition (and tacitly assuming that the relevant feasible end time points *T* lie in a connected set), the minimal value for *T* can be found using any one-dimensional root finding algorithm. For its simplicity and guaranteed convergence, we propose to use the following *bisection method* for the determination of *T* in TOA:

**Input:**

- an initial interval [*t*_min_, *t*_max_] ⊆ [*t*_0_, *t*_end_] for the optimal end time point *T*
- a tolerance level *tol*

**While** |*t*_max_ – *t*_min_| > *tol*:

Set *t*_new_ – (*t*_min_ + *t*_max_)/2
**If** (3.8b), (3.8c) is soluble for *T* = *t*_new_:

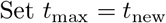
**Else:**

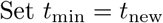

**Output:**

*T* = *t*_new_ as approximation to the optimal end time point

For the initial time interval [*t*_min_, *t*_max_], one needs to assume that (3.8b) and (3.8c) define an infeasible problem on [*t*_0_, *t*_min_], while the corresponding problem on [*t*_0_, *t*_max_] is feasible. The quick convergence of the bisection method entails that an already very good initial guess is not crucial for an efficient implementation, as long as the simulation task is not too computationally expensive.

If there is legitimate doubt about the result, the algorithm can be re-started with another initial interval or one can change to a more fine-grained sampling for the evaluation of feasible and infeasible points. The numerical results in Section 4 were preceded by an exhaustive scan of end time points, which indicated that the set of feasible end time points do indeed form a single connected interval in all shown examples.

#### Remark 3.9.

For the solution of the Pareto problem (3.10), it is not necessary to implement algorithms for maximizing *α*, i.e., optimizing the second objective. Instead, one can continue using the algorithm for time-optimal adaptation while simultaneously fixing feasible values of *α*. With respect to the definition of Pareto-optimality, this means that for any feasible value of one objective, the other one is optimized, corresponding to the so-called *ϵ-constraint method* in multi-objective optimization, cf. [4].

### 3.6 Time-optimal Adaptation Variability Analysis (TOA-VA)

Minimizing *T* need not suffice to uniquely determine the time courses in ***y***. If this is the case, the variability over time can be captured by enumerating possible time series once the optimal end time point was found. We will refer to this procedure as *TOA-Variability Analysis* (TOA-VA). In contrast to static flux variability analysis (FVA), there are several ways to define what such enumeration means. One way would be to determine the maximum and minimum possible value for all components of ***y*** and separately at each time point. This, however, would not only lead to time-consuming computations, but would also be difficult to interpret: a numerical solution that is constructed via putting together maxima or minima 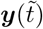 for all time instances 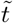 does not have to fulfill the dynamics defined by the original model. Here, we understand TOA-VA as the minimization and maximization of the integral over all components of ***y***, i.e., for all *i* = 1, 2,…, *n_**y**_*:

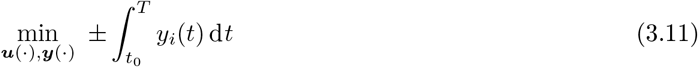

subject to the dynamic and/or boundary constraints in the original problem. Note that the overall time courses might still not be uniquely defined from (3.11).

To explore the variability of the time courses for the concentrations ***c***(*t*), we use the following variant of TOA-VA, which is called *relative TOA-VA*:

i. Compute the optimal end time point *T* of time-optimal adaptation.
ii. For all *i* = 1, 2,…, *n_**y**_*: Use TOA-VA as in (3.11) to obtain a minimal value *I*_min,*i*_ and a maximal value *I*_max,*i*_ for the integral of *y_i_* over [*t*_0_, *T*].
iii. Calculate for all *i* = 1, 2,…, *n_**y**_* the (maximal and minimal) concentrations *c_i_*(*t*) as given by (3.1) where ***y***(*t*) is calculated from

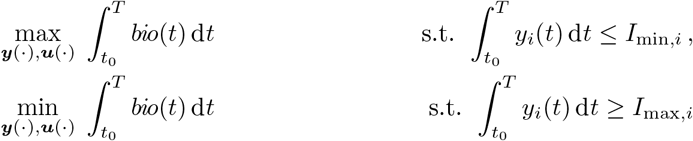

(again subject to the original constraints of the problem).

There is still no guarantee that the solutions to this problem are unique. However, since the concentrations are defined as the ratio of total amounts to the biomass, the above definition is reasonable as one is maximized whilst minimizing the other. Note, that this definition implies that the weighted some of all (maximal or minimal) concentrations no longer needs to add to the total biomass.

### 3.7 Implementation

The calculations for all experiments in Section 4 were done in Python 3.8.1 on a laptop computer. The numerical solutions were determined from a complete parameterization (using the trapezoidal rule) of the compounds and fluxes over the entire time range of interest using *n* = 100 steps on an equidistant grid. This leads to a sparse LP problem which was solved using `gurobipy` on Gurobi 9.0.1 solver [Gurobi, 2021] with standard settings (concerning problem formulation and tolerances). Most experiemnts were repeated (for verification) with tight error tolerances (`‘FeasibilityTol’: 1.0e-9, ‘MarkowitzTol’: 0.999, ‘NumericFocus’: 3, ‘OptimalityTol’: 1.0e-9`) without notable differences. For time-optimal adaptation, no objective vector for the LPs is necessary, so we used the null vector **0**. For (relative) TOA-VA, the integrals in the objective or constraints were approximated using the same time grid and also the trapezoidal rule.

## 4 Results

### 4.1 A simple self-replicator model

We illustrate TOA by means of a coarse-grained self-replicator model [20, 6]. The model, cf. Figure 1, consists of three compounds: *M* (intracellular metabolic precursor), *Tr* (transporter), and *R* (ribosome), as well as five biochemical reactions, together with one external nutrient *N*. The uptake of the external nutrient *N* is catalyzed by the transporter *Tr* and depends on the concentration of *N* via a Michaelis-Menten rate equation. The synthesis of the catalytic macromolecules *Tr* and *R* is limited by the ribosome amount. Within the model, macromolecules can be disassembled into the precursor *M*. For energetic consistency, however, disassembly results in fewer precursor molecules than required for synthesis, reflecting the energy expenditure of protein synthesis and thereby avoiding futile cycles. The metabolic precursor *M* can accumulate and its amount is not required to be balanced at all times, hence *M* also serves as a storage compound. All constraints of the model can be formulated in terms of linear inequalities. A detailed definition is provided in Appendix B.1.

**Figure 1:**
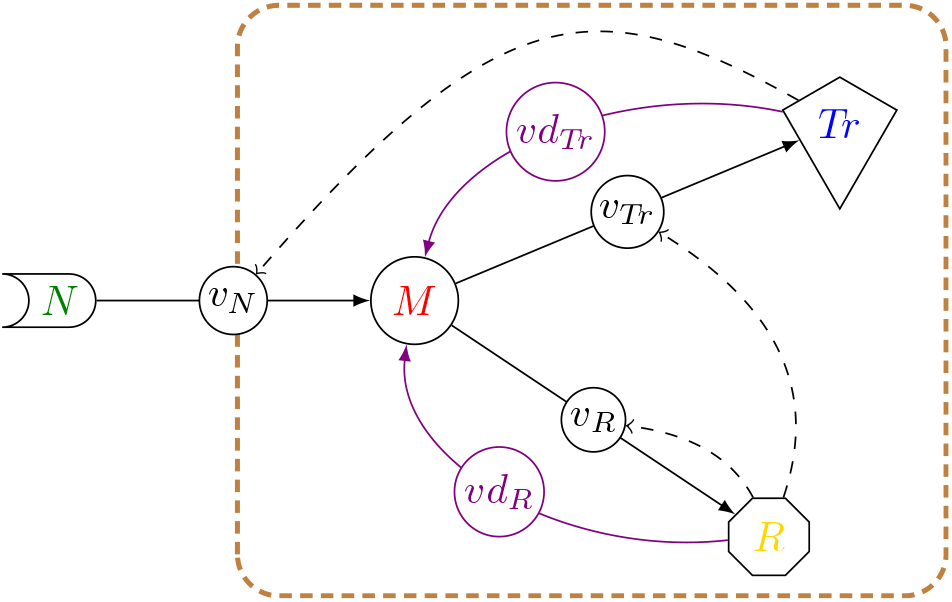
Schematic illustration of the self-replicator model; solid lines represent biochemical reactions between the nodes (biochemical compounds), dashed black lines indicate that a reaction is catalyzed by the respective compound, the brown dashed line symbolizes the cell’s boundary. Abbreviations: *N*: external nutrient, *M*: metabolic precursor/storage, *Tr*: transporter, *R*: ribosome

### 4.2 Perfectly adapted cells and RBA

Before the dynamic behavior of the model is studied by means of TOA, we summarize basic properties of the model in a constant environment using Resource Balance Analysis (RBA). RBA provides a way to calculate the steady-state amounts of the cell maximizing the total cell growth under constant external conditions.

Figure 2A shows the maximal growth rate λ as a function of the relative nutrient availability. The growth rate follows a Monod equation with a maximum λ^*max*^ ≈ 0.435 h^-1^ and an effective (dimensionless) affinity constant *K_A_* ≈ 0.347, corresponding to the value of the relative nutrient availability *N/K_M_* at which the cell grows at half the maximal growth rate λ^*max*^.

**Figure 2:**
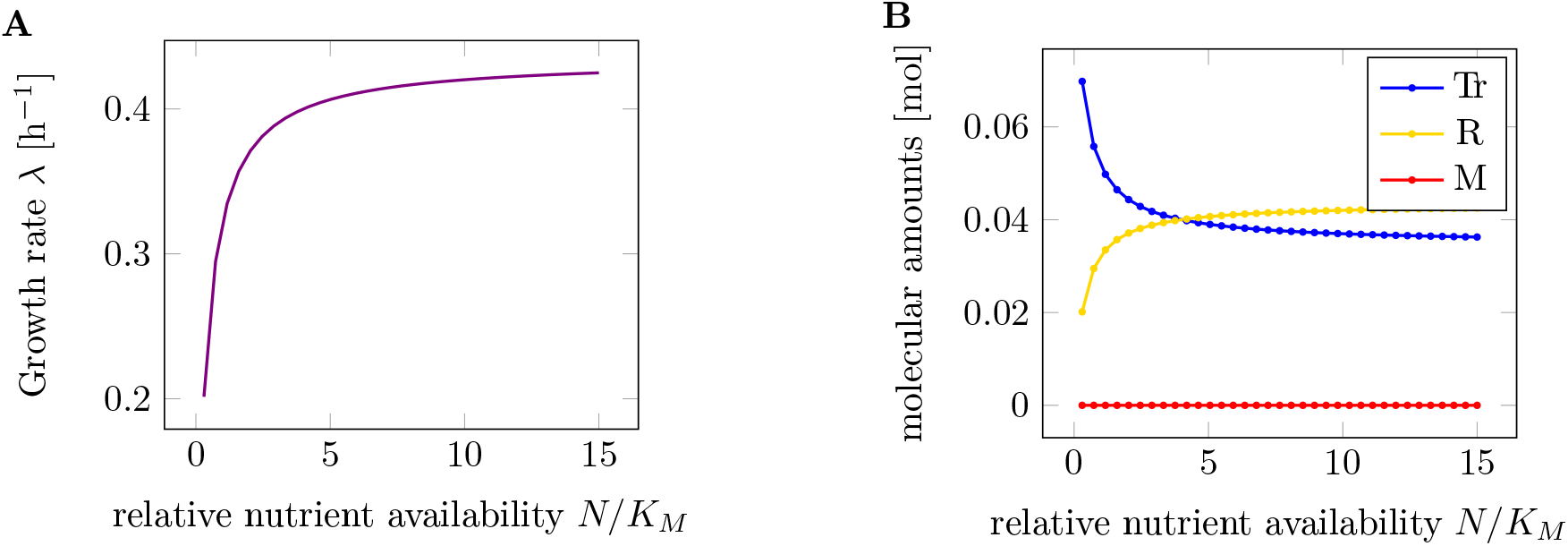
**A**: Maximal growth rate λ as a function of extracellular nutrient availability as predicted by RBA. **B**: Cellular amounts of intracellular compounds as functions of relative nutrient availability. The latter is scaled with respect to the Michaelis constant *K_M_* = 1.0 mol of the uptake reaction.

Figure 2B shows the total amounts of the three intracellular components *M*, *Tr*, and *R* as a function of the (relative) nutrient availability. The amounts were scaled such that the total biomass always equals one unit (e.g. 1 g cellular dry weight). As expected, when maximizing the growth rate, the optimal level of the precursor/storage component *M* is always zero. This reflects the fact that the precursor *M* has no catalytic activity, and any non-zero amount of *M* would consume resources that otherwise could be allocated to transport or protein translation.

The amounts of the other intracellular components *Tr* and *R* follow the well-known *growth laws* of microbiology [27]. The concentrations are a function of the growth rate, and hence the external nutrient availability, the well-known linear relationship is shown in Appendix E. With increasing nutrient availability, the relative amount of transporter decreases, whereas the relative amount of ribosome increases.

### 4.3 TOA in constant environments

Our first case study using TOA is to consider the doubling of a microbial cell in minimal time. We assume that the relf-replicator model in Figure 1 has pre-described initial amounts ***y***(*t*_0_) = ***y***_0_ and the objective is to double all cellular components as fast as possible, cf. Application 3.1. The environment is assumed to be constant with an relative (external) nutrient availability *N/K_M_* = 1. The initial (and final) amounts are not assumed to be optimal for the given environment. Instead, ***y***(*t*_0_) is obtained by solving an RBA problem corresponding to *N/K_M_* = 2.0. In other words, the cell is assumed to be adapted to a higher nutrient level than is present in the current environment. In the following, we will refer to such cells as “*optimists*”.

Figure 3 shows the time course of intracellular components for one cell doubling. The predicted time-optimal amounts of metabolic compounds are shown as solid lines (red, blue, and yellow), the total biomass is shown in green. The dashed lines correspond to a solution obtained by iterative RBA (cf. Example 3.4), which corresponds to exponential growth of all cellular components with no further internal degrees of freedom. Figure 3B shows the respective flux rates over the simulated time range. Solid lines again indicate the solution of TOA, while dashed lines (exponential curves) correspond to the solution found with iterative RBA.

**Figure 3:**
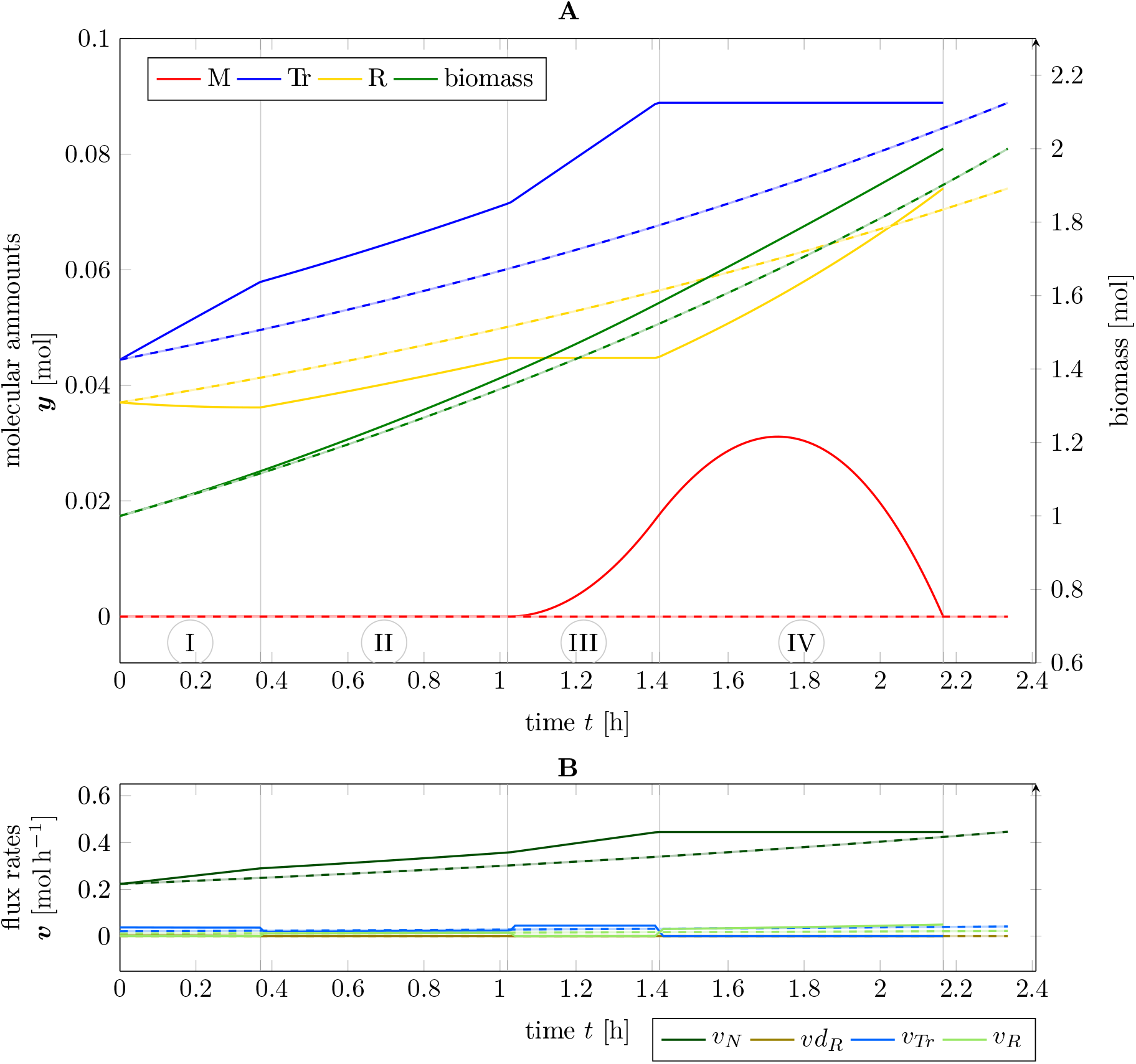
Cell cycle of an “optimistic” cell; **A:** amounts and biomass as a function of time, **B:** flux rates as a function of time; solid lines indicate the solution of TOA, dashed lines indicate iterative RBA (exponential growth) with the same “optimistic” initial values.

Using TOA, the time for one cellular doubling is *T* = 2.17 h. In contrast, the solution based on iterative RBA results in a slightly longer doubling time of *T* = 2.34 h, showing that internal degrees of freedom shorten the calculated division time. The time course of ***y***(*t*) over one cell doubling can be subdivided into four time intervals (marked as I-IV in Figure 3A). At the beginning (marked as interval ‘I’), cell growth is limited by the lack of transporter *Tr* due to the “optimistic” initial configuration of the cell. Hence, ribosome *R* is actively disassembled into precursor *M* to increase the synthesis of *Tr*. In interval ‘II’, the cell is perfectly adapted to the given nutrient environment and grows exponentially, before the re-adaptation to the target composition ***y***^goal^ = 2 · ***y***^init^ begins in interval ‘III’. Within interval ‘III’, the cell still has an overabundance of *Tr*, which allows it to accumulate the precursor *M*. In the final interval ‘IV’, synthesis of transporter *Tr* ceases and all resources are devoted to the synthesis of the ribosome *R*, until the target amounts ***y***^goal^ are reached.

The biological plausibility of these time courses is discussed in Section 5. Here we only summarize the following results: Given the initial amounts ***y***^init^, cell doubling using TOA in time-invariant environments gives rise to complex intracellular dynamics different from solutions obtained by iterative RBA. Importantly, these solutions involve a transient accumulation of the precursor *M* as a storage compound-a phenomenon not observed with iterative RBA. The minimal division time predicted by time-optimal adaptation is shorter than division times obtained by iterative RBA.

So far, we considered a particular initial amount ***y***^init^ such that the cell was adapted to a higher nutrient availability than actually present in the environment (“optimist”). To obtain a broader view, we evaluated cell doubling using TOA in different time-invariant environments with initial (and final) amounts adapted to different external nutrient availability. The results are shown in Figure 4. Solid lines correspond to intracellular amounts using TOA, dashed lines correspond to a solution obtained with iterative RBA (exponential growth without internal degrees of freedom). Shaded areas correspond to variability in the sense of TOA-VA (cf. Section 3.6), i.e., possible solutions that equally satisfy all constraints and the optimality criterion. In this case, the solid lines display a ‘nominal’ solution, i.e., one that was provided by the algorithm before an additional variability analysis (we note that since the numerical solution is based on a feasibility problem, the LP solver has no incentive to favor a smooth solution to any other).

**Figure 4:**
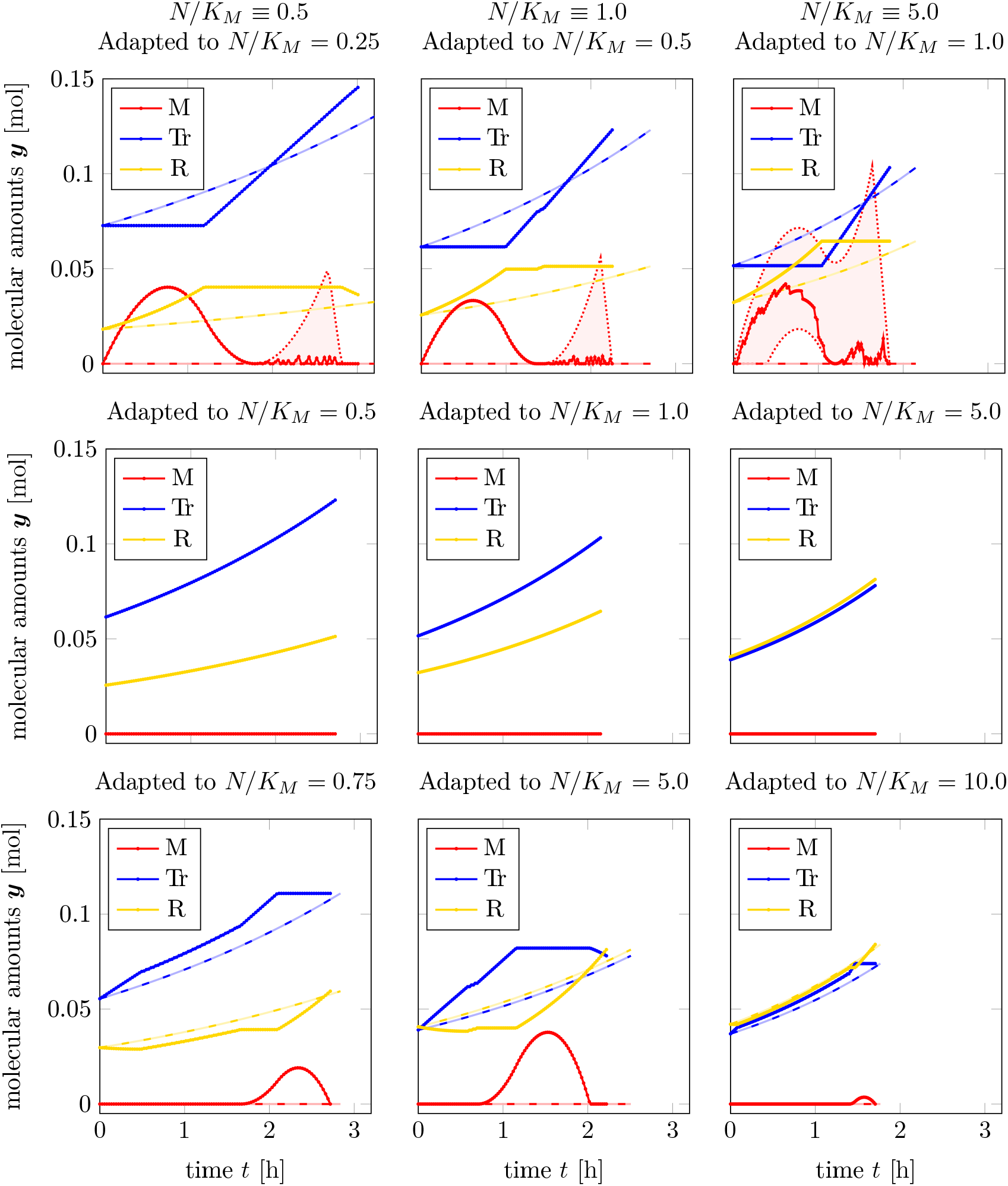
Time course solutions of time-optimal adaptation and a cell doubling experiment under different constant external nutrient conditions; solid lines: TOA, shaded areas: TOA-VA, dashed lines: iterative RBA (simulated until cell doubling was achieved); upper row: pessimistically adapted, middle row: perfectly adapted (recovery of iterative RBA), bottom row: optimistically adapted for constant relative nutrient availability of *N/K_M_* = 0.5 (left column), *N/K_M_* = 1.0 (middle column), and *N/K_M_* = 5.0 (right column)

Columns in Figure 4 correspond to different relative nutrient availability levels: the first column to a nutrient availability *N*(*t*)/*K_M_* ≡ 0.5; the second column to *N*(*t*)/*K_M_* ≡ 1.0, and the third to *N*(*t*)/*K_M_* ≡ 5.0. The rows in Figure 4 correspond to different “expectations” of the cells, that is, which external nutrient availability the initial (and final) amounts are adapted to. Specifically, the first row corresponds to “pessimists”. That is, cells adapted to a nutrient availability below the one present in the environment, while retaining the objective to double all cellular components in minimal time. The second row corresponds to cells perfectly adapted to the environmental nutrient availability. The final row corresponds to “optimists”, i.e., cells adapted to a higher nutrient availability than present in the environment.

The latter scenario corresponds to the example already shown in Figure 3. We again observe an initial increase in the transporter synthesis, followed by a delayed onset of ribosome synthesis. Importantly, in each case, we can see a transient accumulation of storage *M*(*t*) that is absent in solutions obtained by iterative RBA. In the case of perfectly adapted cells (middle row), solutions obtained by TOA are equivalent to solutions obtained by iterative RBA. For “pessimistic” cells (top row), we again observe complex time courses. In particular, cells adapted to lower nutrient levels than present in the environment exhibit an overabundance of transporter. Hence, we observe an initial rapid uptake of nutrient and transient accumulation of the precursor *M*. In the initial interval, resources are primarily allocated to the synthesis of ribosomes. Only in the later interval, the transporter is synthesized to the required amounts (even at the expense of ribosomes that may be disassembled into precursors). The transient accumulation of precursor *M* exhibits considerable variability and the solutions of TOA are no longer unique.

The biological plausibility of these time courses is again discussed in Section 5. Here we only note that, despite the simplicity of the model, the solutions exhibit a wide variety of qualitatively different complex temporal behaviors, including the transient accumulation of the precursor *M*.

### 4.4 The role of expectation: optimists versus pessimists

We further investigate two key observations obtained in the previous experiments: the transient accumulation of precursor *M* as a storage compound, as well as the impact of the initial cellular state on the predicted doubling time.

Firstly, Figure 5 shows the average storage concentration predicted for a population of cells adapted to a different nutrient availability (*N/K_M_* ∈ (0.2, 2.0), x-axis) in an environment with an actual relative nutrient availability *N/K_M_* ≡ 1.0. To calculate the average storage concentration predicted by TOA for a population of cells, we assume that the (*in silico*) measurements are taken from a heterogeneous population of unsynchronized cells that are (equidistributed) at various stages of a cellular doubling. To take this non-uniform age distribution into account, the population average was computed, cf. [23], as

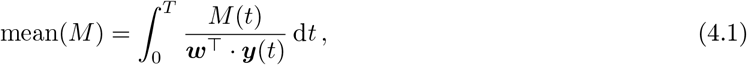

where ***y***(*t*) is a solution obtained by *relative TOA-VA*, cf. Section 3.6.

**Figure 5:**
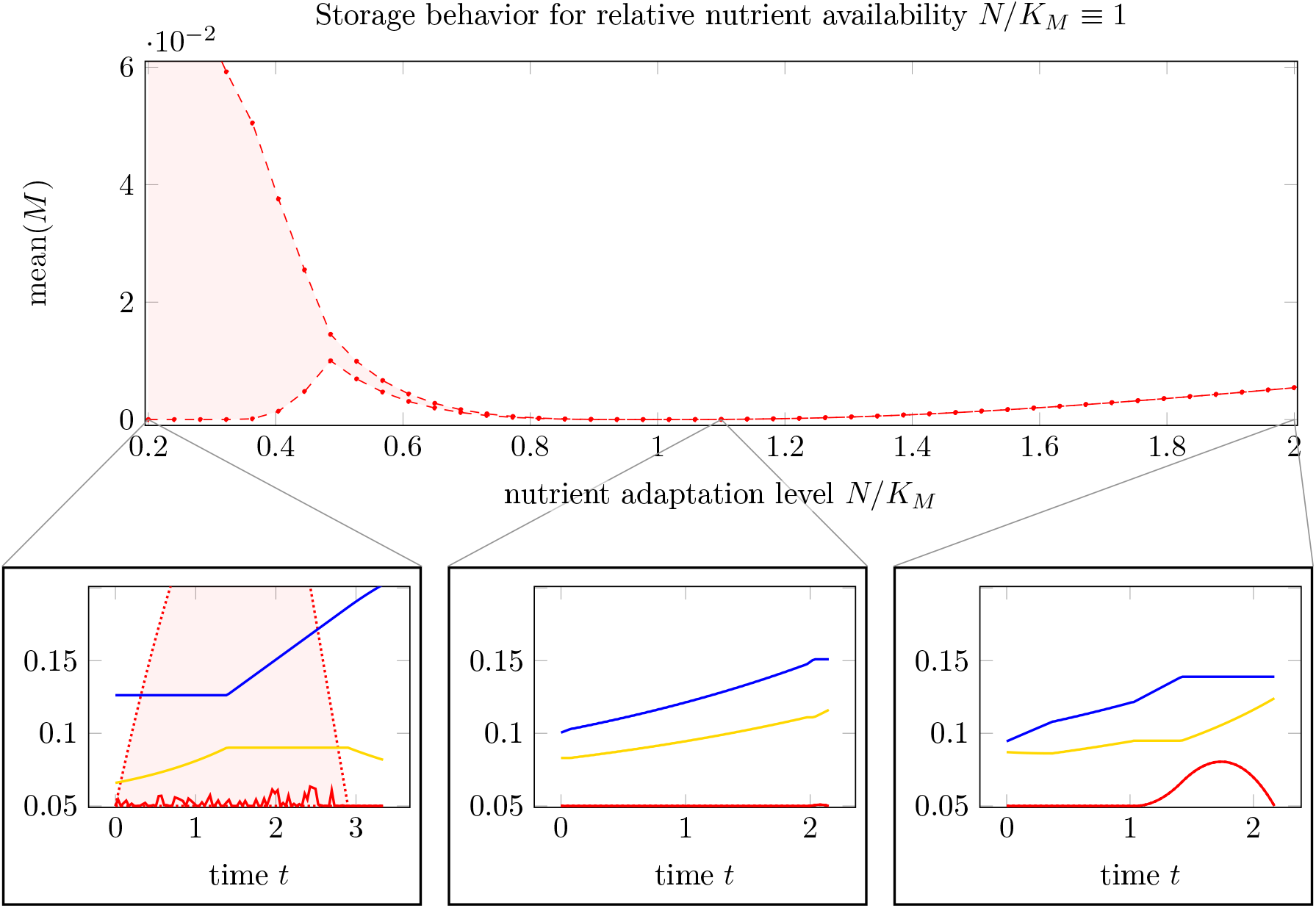
Influence of optimistic and pessimistic goal states in cell doubling: Main plot: mean relative storage accumulation, see (4.1), as a function of nutrient adaptation level. Bottom row: Three selected time courses, cf. Figure 4, for nutrient adaptation levels *N/K_M_* of 0.2, 1.1, and 2.0. For *N/K_M_* < 1, the quantity mean(*M*) is no longer unique such that a shaded area indicates the possible range, as TOA-VA also predicts a range of possible solutions (shaded area in the bottom left plot).

As shown in Figure 5, we observe (the possibility of) a *nonzero* average storage concentration for all cellular states that are not perfectly adapted to the respective environment. For optimistic cells adapted to a higher nutrient availability than present in the environment, the average storage concentration increases slightly with the distance to the perfectly adapted state. The effect is more pronounced for pessimistic cells adapted to a lower nutrient availability than present in the environment. In this case, the solutions of TOA are not unique and the *range* of average storage is indicated as a shaded area. For “pessimist” cells, the large average storage is due to a high abundance of transporter molecules, which implies that uptake and accumulation of precursor is not restricted.

Secondly, Figure 6 shows the predicted growth rate for cells adapted to a different relative nutrient availability (*N/K_M_* ∈ (0.2, 2.0)) than present in the environment (*N/K_M_* ≡ 1.0). The straight line indicates the growth rate of cells that are perfectly adapted, resulting in a maximal growth rate λ = λ_env_ ≈ 0.32 h^-1^. The maximal growth rates for cells adapted to a different environment (misadaptation) are shown as a solid green line for solutions obtained with TOA and as a purple line for solutions obtained with iterative RBA.

**Figure 6:**
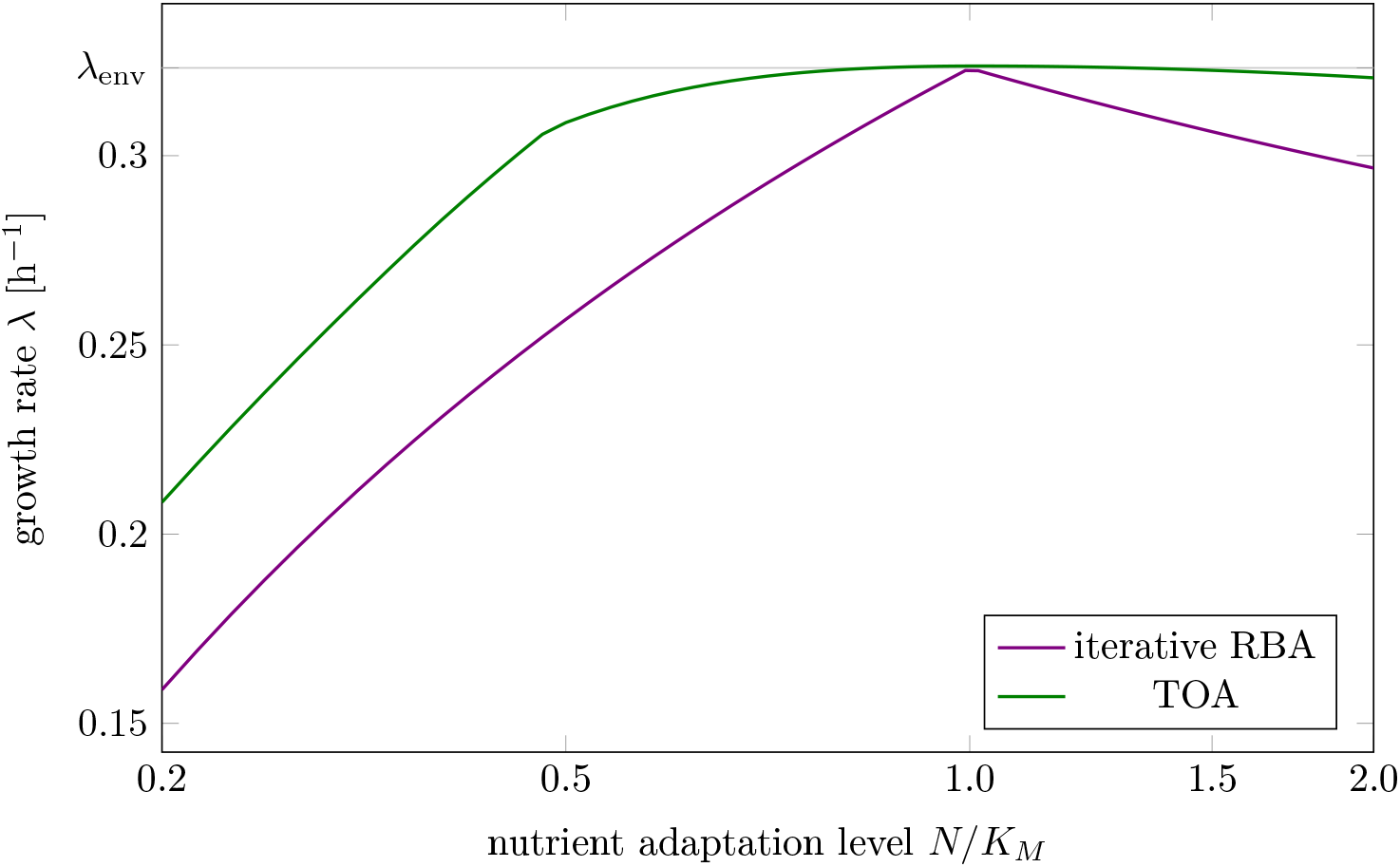
Growth rate of differently adapted cells as predicted by cell-doubling experiments using TOA and iterative RBA in an environment with relative nutrient availability *N/K_M_* = 1.0; λ_env_ ≈ 0.32 h^-1^ denotes the maximal growth rate as predicted by RBA.

We observe that misadaptation always results in a reduced growth rate, as compared to a perfectly adapted cell. However, solutions obtained by TOA always outperform solutions obtained by iterative RBA, demonstrating that internal degress of freedom and transient accumulation of storage shorten the predicted doubling time. Furthermore, the decrease in growth rate is more pronounced for “pessimistic” adaptation, that is, for cells that are adapted to a lower nutrient level than present in the environment. In contrast,”optimistic” adaptation, that is, cells are adapted to a higher levels than present in the environment, together with TOA results in growth rates close to perfectly adapted cells-indicating that “optimistic” adaptation carries a lower evolutionary cost than “pessimistic” adaptation.

### 4.5 Time-optimal adaptation at a nutrient shift

As our second application, we consider a nutrient shift, i.e., a sudden change in the external conditions from a given constant nutrient availability for *t* < 0 to a different one for *t* ≥ 0. TOA is utilized to predict the time-optimal transition of a cell perfectly adapted to the initial state at *t* < 0 to a state perfectly adapted to maximize growth in the new environment for *t* ≥ 0. As noted in Section 3.4, the target state for the new environment is typically defined in terms of concentrations rather than amounts, because it is unknown whether or how much the cells are able to grow during adaptation.

Figure 7 shows the resulting time courses for the coarse-grained self-replicator model used in the previous sections. Shown are time-optimal shifts from a low nutrient availability to a higher nutrient availability (left column in Figure 7), as well as time-optimal shifts from a high nutrient availability to a lower nutrient availability (right column in Figure 7). Non-unique solutions are again displayed as shaded areas indicating the maximum and minimum range in which solutions can be found (TOA-FVA, see Section 3.6). We observe that the time-optimal transition from lower to higher nutrient availability again entails a transient accumulation of storage.

**Figure 7:**
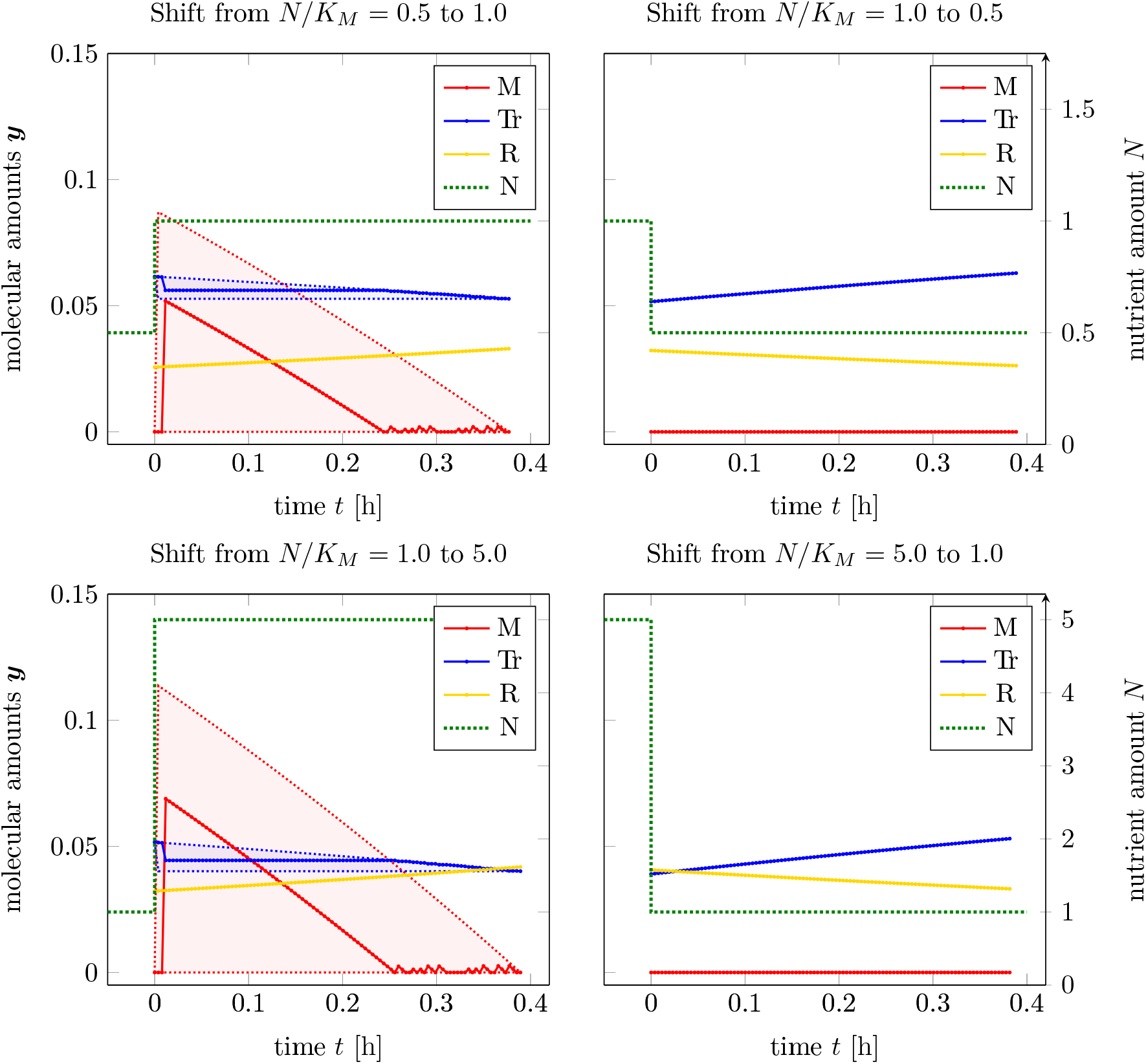
Adaptation to a single nutrient jump (shown as a dashed green line), left column: adaptations from poorer to richer medium, right column: adaptation to scarcer environment; shaded areas: solutions in the sense of TOA-VA. We note that for *t* < 0, TOA makes no assumptions about ***y***(*t*).

As detailed in Section 3.4, time-optimal adaptation alone may fall short as an evolutionary principle to explain cellular adaptation after a nutrient shift. Rather, we consider a two-objective optimization in the sense of Pareto with the conflicting objectives of a fastest possible adaptation to the new state versus a maximal increase in total cellular biomass.

Figure 8 (main panel) shows the resulting Pareto fronts for different transitions in terms of the minimal time *T*^*^ for adaptation versus the maximal increase in cellular biomass given by the factor *α*, cf. (3.10). Panels A-D in Figure 8 show selected time courses of intracellular amounts at different locations of the Pareto front. In the subplots A and B, the shaded areas indicate when the cell is perfectly adapted to the environment in the sense of RBA, i.e., from the start of the shaded areas, the cell is exponentially growing at a maximum possible growth rate and no further internal dynamics take place. The absence of internal dynamics explains that, for larger values of *α* or *T*\*, the lines in the main plot become asymptotically parallel.

**Figure 8:**
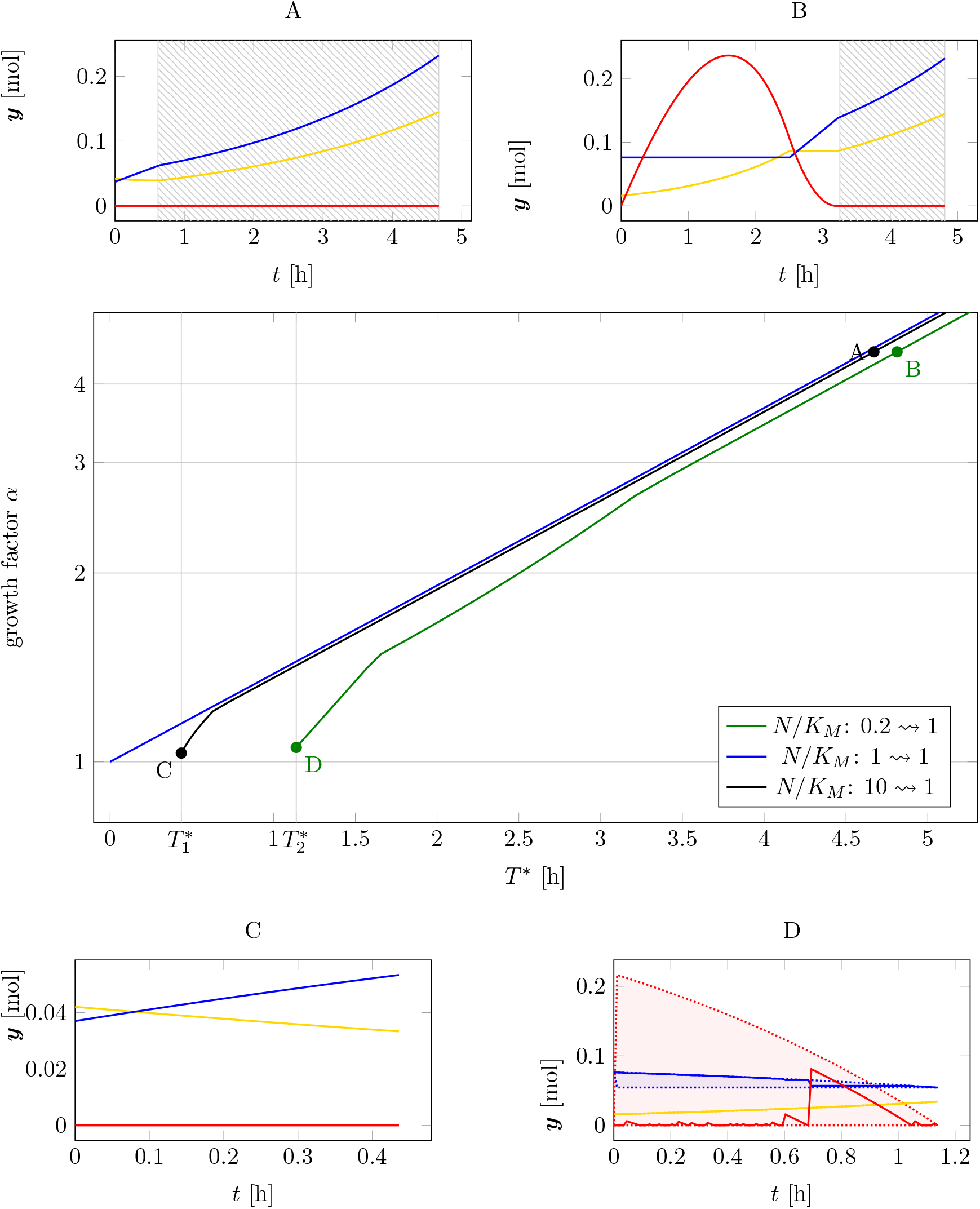
Two-objective optimization of adaption time *T*\* and total biomass growth factor α for time-optimal adaptation at a nutrient shift. Main plot: Pareto fronts for three different initial adaptations (measured in *N/K_M_*) of 0.2, 1.0 and 10. Subplots A-D: Time courses at different points on the Pareto fronts, cf. Figure 7. The shaded areas in subplots A and B indicate time intervals where the cell is perfectly adapted (exponential growth).

In the absence of a nutrient shift (i.e., the transition *N/K_M_*: 1 → 1, blue line in Figure 8), the minimal time for adaptation is *T*^*^ = 0 with a growth factor *α* = 1, and the relationship between transition times *T*^*^ > 0 and increase in biomass is consistent with exponential growth (note the logarithmic scale on the y-axis).

For a nutrient shift from high to low nutrient availability (*N/K_M_*: 10 → 1, black line) the minimal transition time is 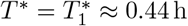. Figure 8A and C show two representative transitions on the Pareto front with panel A corresponding to a scenario that prioritizes an increase in biomass (factor α) over the transition time *T*^*^, and panel C a scenario that prioritizes a minimal transition time over the accumulation of biomass.

For a nutrient shift from low to higher nutrient availability (*N/K_M_*: 0.2 → 1, green line) the minimal transition time is 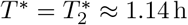. Figure 8B and D show two representative transitions on the Pareto front with panel B corresponding to a scenario that prioritizes an increase in biomass (factor *α*) over the transition time *T*^*^, and panel D a scenario that prioritizes a minimal transition time over the accumulation of biomass. In either case, the optimal transition involves a transient accumulation of the storage compound *M*.

Consistent with results in the previous section, Figure 8 also shows that “optimistic” adaptation carries a lower evolutionary cost than “pessimistic” adaptation. A cell adapted to high nutrient availability exhibits only a slightly reduced biomass increase when transitioning into a low nutrient environment, as compared to a cell already adapted to this environment. In contrast, a cell adapted to a lower nutrient environment exhibits a more pronounced reduction in accumulated biomass when transitioning into higher nutrient availability, as compared to either a cell that is already adapted to the higher nutrient availability, or likewise as compared to a cell that was previously adapted to even higher nutrient availability.

## 5 Discussion

In this work, we introduced TOA, a novel approach to simulate and predict time-optimal adaptation of microbrial metabolism and growth. While time-optimal modeling has been considered before, see, among others, [1] (minimization of lag/response-time), [30] (maximize survival time under nutrient depletion), [3] (bio-reactor applications), or [14] (temporal gene expression), our work builds upon the recent advances in dynamic constraint-based modeling, such as dFBA, deFBA and cFBA, cf. Section 3.2. Our method, TOA, is versatile and extends most approaches currently employed in constraint-based modeling of microbial metabolism and growth.

We exemplified the use of TOA by considering two prototypical applications, the doubling of a cell in a constant environment (cf. Application 3.1), as well as the time-optimal adaptation to a nutrient shift (cf. Application 3.2). The application of TOA was illustrated by means of a coarse-grained self-replicator model, and the results illustrate the utility of TOA to generate and explore biological hypotheses.

Specifically, the premise underlying our *in silico* experiments for cell doubling in a constant environment was that microbial cells are not necessarily precisely adapted to the given environment, but may nonetheless have evolved a regulatory scheme that allows them to double their intracellular composition in minimal time. Based on this premise, TOA predicts (i) complex intracellular dynamics different from solutions obtained by iterative RBA, (ii) that transient accumulation of storage compounds reduces the predicted doubling time, and (iii) that (mis-)adaptation to a higher nutrient availability than actually present in the environment carries a lower evolutionary cost than (mis-)adaptation to a lower nutrient availability.

While, due to the simplicity of the coarse-grained model, we do not expect the specific time courses obtained for the model to be exact predictions of biological reality, we are confident that the results reveal several genuine biological insights. In particular, the role of storage compounds in cellular metabolism is not fully understood. Beyond the role of storage in diurnal oscillations, cf. [26, 25] and as a safeguard for periods of nutrient scarcity, our simulations show that *even under constant environmental conditions*, storage may play an important role. Under certain conditions, intracellular dynamics and transient accumulation of nutrients can indeed reduce doubling time–a fact that is of potential relevance considering that cells indeed exhibit coordinated metabolic dynamics over a cell cycle [22].

These results are further supported by the simulation of the time-optimal cellular adaptation to a nutrient shift. While it has been shown that storage compounds, such as glycogen, provides short-term benefits in changing environments [28], most current constraint-based methods do not predict the accumulation of storage compounds. In contrast, TOA shows that transient accumulation of storage can reduce the time required for adaptation. In particular, the rapid uptake and storage of nutrients following an upshift in supply (as shown in Figure 7, left column) is reminiscient of “luxury uptake” or “over-compensation”. The latter phenomenon occurs when cells are starved and re-exposed to a limiting nutrient, such as phosphate, and can be exploited, for example, for nutrient removal from wastewater [24]. While the molecular details of such phenomena are not well understood, our analysis using TOA shows that such “over-compensation” or “overshoot” phenomena can indeed be explained using principles of (optimal) cellular resource allocation-and do not necessarily require explanations that invoke competition between individuals to rationalize rapid nutrient uptake after starvation.

Finally, the results of TOA show that the costs of mis-adaptation to an environment are not symmetric, neither for cell doubling in a constant environment (Figure 6), nor for adaptation after a nutrient shift (Figure 8). In either case, a cell that is adapted to a higher level of (extracellular) nutrient than available in the environment (“optimist”) has only a minor disadvantage compared to an already perfectly adapted cell. Vice versa, however, cells that are adapted to a lower level of (extracellular) nutrient than available in the environment (“pessimist”) have a pronounced disadvantage compared to a perfectly adapted cell. This asymmetry indicates that adaptation to a low nutrient environment is only advantageous if the low nutrient state persists for extended periods of time. And indeed, it has been suggested that some microorganisms, such as *Lactococcus lactis*, may simply preserve a large overcapacity of ribosomes and glycolytic enzymes to be ready to rapidly respond and grow when conditions improve, and thereby implement an “optimistic strategy” [7].

## 6 Conclusions and Outlook

Constraint-based optimization plays an important role to elucidate and eventually predict cellular behavior. As an extension of previous modeling frameworks, we introduced time-optimal adaptation. TOA is motivated by the assumption that under certain conditions it is evolutionary favorable to adapt to a new cellular state in minimal time. In its general form, TOA can be applied in a very broad sense and thereby extends most of the existing constraint-based modeling frameworks.

As shown in this work, TOA allowed us to elucidate several biological phenomena, such as the accumulation of storage in constant environments and “overshoot” accumulation of nutrients after starvation, which cannot be readily simulated using existing methods-thereby demonstrating the utility of TOA for future analysis.

While the examples discussed within this work focused on constant environments and simple nutrient shifts, TOA can also be applied in time-dependent environments and can be readily extended to include further constraints. Likewise, as shown in this work, TOA can be included within multi-objective optimization in the sense of Pareto.

Possible further extensions include “t-max adaptation”, i.e., to maximize, for example, survival time under nutrient starvation, as well as more general constraints on the target state (for example, to attain a minimal amount of a specific intermediate in minimal time, while the amounts other cellular components are not specified).

We are therefore confident that TOA and its possible extensions are a valuable contribution in the context of constraint-based modeling with manifold applications beyond the examples discussed in this work.

## A Abbreviations and Notations

***y***(*t*): total molecular amounts (of metabolites) within the cell
***v***(*t*): flux rates in the metabolic network
S: stoichiometric matrix
***u***(*t*): collection of (mostly time-dependent) degrees-of-freedom in the network (mostly flux rates)
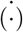: time derivative, e.g. 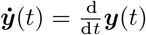
***w***: vector of molecular weights
*bio*(*t*): biomass ***w***^┬^ · ***y***(*t*) of the cell
***c***(*t*): concentrations of metabolites 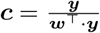
0, 0: zero vector, zero matrix
[*t*_0_, *T*^*^]: time interval of interest
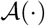: (abstract) set defining the constraint-based modeling framework, see (3.8c)
λ: Growth rate of the cell (given exponential growth from mass *m*_0_ to mass *m*_1_ in a time interval of length 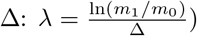
(·)^┬^: transpose of a vector/matrix
(·)_*i*_1_:*i*_2__: indexing of a vector: For ***a*** = (*a*_1_, *a*_2_,…, *a_n_*)^┬^ and positive integers *i*_1_ ≤ *i*_2_, it holds ***a***_*i*_1_:*i*_2__:= (*a*_*i*_1__, *a*_i_1_+1_,…, *a*_i_2__)^┬^. If no start/end index is supplied, all remaining indices are included: ***a***_*i*_1:__:= (*a*_*i*_1__, *a*_*i*_1_ + 1_,…, *a_n_*)^┬^
≤: (component-wise) smaller-or-equal relation
LP: Linear program/linear programming
TOA(-VA): time-optimal adaptation (Variability Analysis)
FBA: flux balance analysis
FVA: flux variability analysis
cFBA: conditional FBA
RBA: resource balance analysis
deFBA: dynamic enzyme-cost FBA

## B Self-Replicator Model

### B.1 Model Description

#### Compounds

The vector of metabolic compounds is given by ***y***(*t*) = (*M*(*t*), *Tr*(*t*), *R*(*t*))^┬^; additionally, there is an external nutrient supply of total amount N(t) that is controlled by the external conditions.

**Reactions:** The vector of reaction fluxes is given by ***u***(*t*) = (*v_N_*(*t*), *v_Tr_*(*t*), *v_R_*(*t*), *vd_Tr_*(*t*), *vd_R_*(*t*))^┬^ with the reactions

- 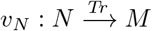
- 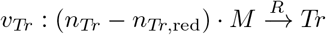
- 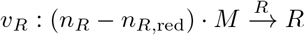
- *vd_Tr_*: *Tr* → *n_Tr_* · M
- *vd_R_*: *R* → *n_R_* · *M*

**Constraints:**

1. Positivity: *M*(*t*), *Tr*(*t*), *R*(*t*) ≥ 0
2. Irreversibility: *v_N_*(*t*), *v_Tr_*(*t*), *v_R_*(*t*), *vd_Tr_*(*t*), *vd_R_*(*t*) ≥ 0
3. Dynamics:

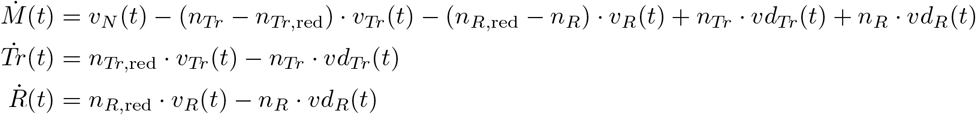
4. Enzyme Capacity constraints:

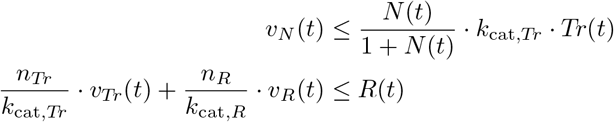

**Constants:**

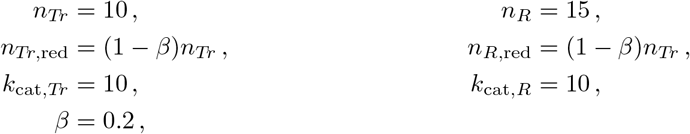

### B.2 Re-formulation as a deFBA problem

In the terms of Example 3.2, the dynamics of the self-replicator model can be described via:

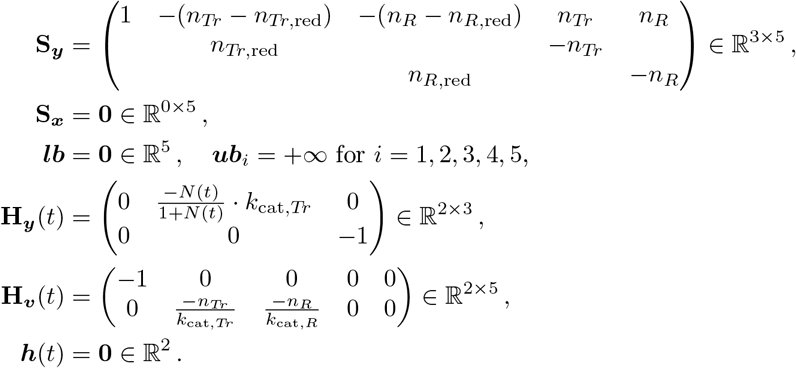

### B.3 Formulation including Extracellular Compound Dynamics

In the above formulation of TOA, extracellular compounds (like nutrients or waste products) are not explicitly incorporated into the framework and all dynamic evolution (by means of ODEs) is restricted to the total cellular amounts ***y***(*t*). The influence of the environment to the cell is captured by the abstract constraint set 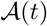 in the general description (3.4) or, more specifically, by the matrices/vectors **H**_***y***_(*t*), **H**_***v***_(*t*), ***h***(*t*) in (3.6). If the amounts of extracellular compounds (now denoted by 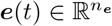) are part of the unknowns in the model, the constraint-based framework (3.4) can be adapted to include dynamic relations, i.e., DAEs including ***ė***(*t*) as well. Formula (3.4a) can then be extended to

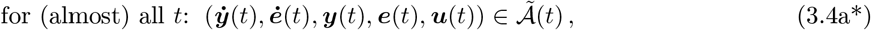

with an appropriate extension 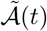 of the original constraint set 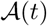. All frameworks covered by (3.4) can be formulated in terms of (3.4a*) as well and, accordingly, TOA can also be formulated with ***e***(*t*) as part of the dynamic variables. However, we note that providing goal states for external biochemicals is more unrealistic from the modeling viewpoint as the cell only has very limited control over its surroundings.

Instead of re-formulating the frameworks in Examples 3.1, 3.2, 3.3 and 3.4 in detail, we will just show here (i) how the self-replicator model can be re-framed to include limited total nutrient availability by considering dynamics of *N*(*t*) and (ii) how the original model can be recovered again by adapting the flux vector. As the uptake reaction is nonlinear (Michaelis-Menten kinetics), a fully linear model is no longer possible.

i. Introduce vectors ***y***(*t*) = (*y_i_*)_*i*=1,2,3_ = (*M*(*t*), *Tr*(*t*), *R*(*t*))^┬^) ≥ **0**, ***e***(*t*) = (*e_i_*)_*i*=1_ = (*N*(*t*)) ≥ 0, ***v*** = (*v_N_*, *v_Tr_*, *v_R_*, *vd_Tr_*, *vd_R_*)^┬^ ≥ **0**, and let

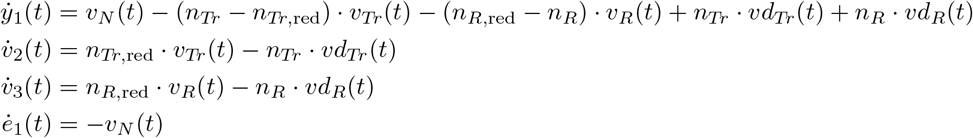

subject to:

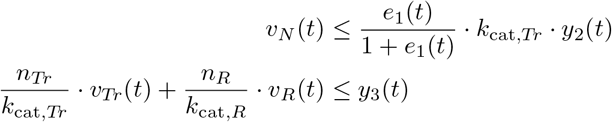

and provide initial values for the external nutrient *N*(*t*_0_). In this formulation, no external nutrient source is present anymore and *N*(*t*) gets accordingly depleted over time (for TOA, this would have important implications as the environment the cell adapts to is no longer the same after the adaptation process).
ii. If the original problem were to be recovered while still including the dynamics of ***e***(*t*), one can (there are multiple ways to model this) introduce another flux rate *v*_in_(*t*) ≥ 0 that compensates the reduction of the nutrient. The differential equation for *e*_1_(*t*) would then need to be adapted to

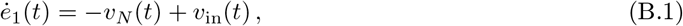

and the overall nutrient level could be kept at a given level *N*_given_(*t*) via (in-) equality constraints

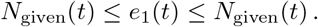 Numerically, this re-formulation is (unnecessarily) complex as the differential equation (B.1) is merely a dummy equation and a linear problem is formulated in nonlinear terms. We state the alternative form here to show that a more general approach using external compounds ***e***(*t*) is able to capture more general biochemical questions; the above restriction to dynamics of ***y***(*t*) was mainly for notational convenience.

## C Formulation in terms of concentrations

The constraints on the boundary values are given by

1. Initial values (as usual):

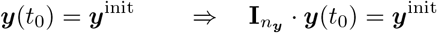
2. Final values:

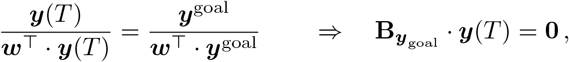

where 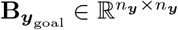 is defined through

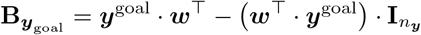

This means, the boundary conditions to be imposed by the optimal control solver are given by

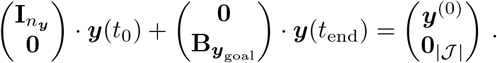

## D Iterative RBA: Characterization as a constraint-based problem

In the notation of the constraint-based framework (3.4), iterative RBA can be written as

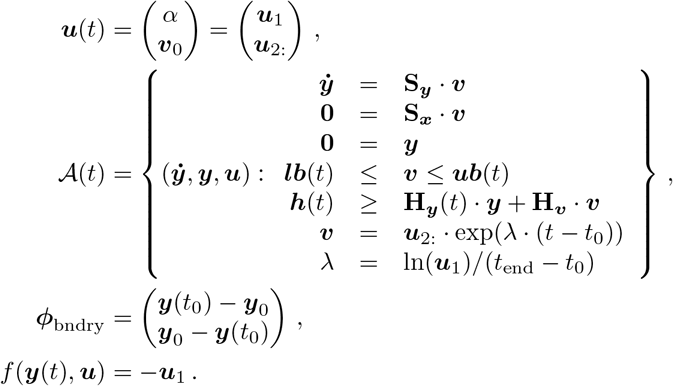

Note that the control variables ***u*** are not time-dependent, i.e., they enter the model as control parameters.

## E Enzyme Fractions relative to Growth Rate

The following plot shows the relation between “optimal” molecular amounts (as predicted by RBA) and growth rate of the cell, cf. Figure 2 and [27].

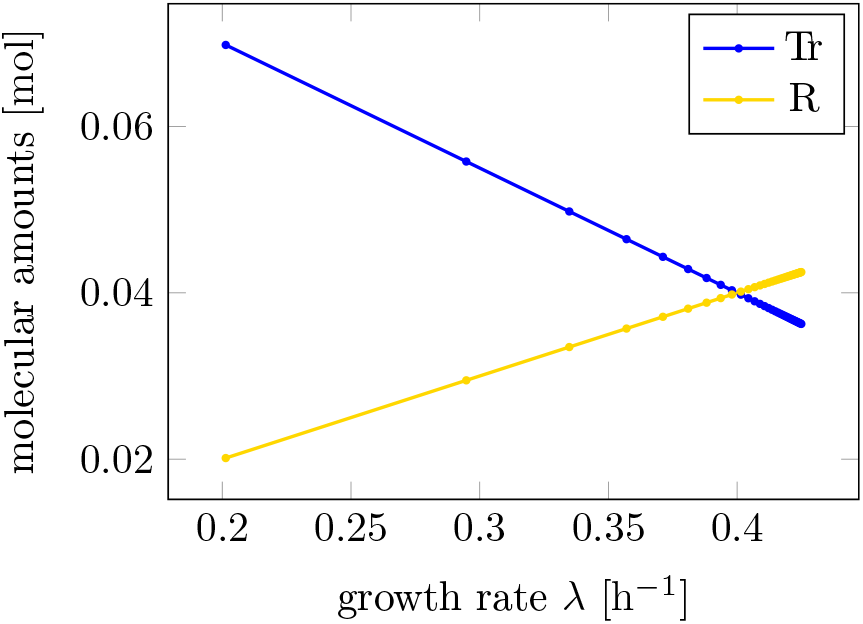

